# Histone deposition plays an active role in recombination-dependent replication to balance genome stability upon replication stress

**DOI:** 10.1101/303305

**Authors:** Julien Hardy, Dingli Dai, Anissia Ait Saada, Ana Teixeira-Silva, Louise Dupoiron, Fatemeh Mojallali, Karine Fréon, Francoise Ochsenbein, Brigitte Hartmann, Sarah Lambert

## Abstract

Replication stress poses a serious threat to genome and epigenome stability. Recombination-Dependent-Replication (RDR) ensures DNA synthesis resumption from arrested forks. Despite the identification of chromatin restoration pathways during DNA repair processes, crosstalk coupling RDR and chromatin assembly is largely unexplored. Here, we addressed the contribution of chromatin assembly to replication stress in fission yeast. We expressed a mutated histone (H3-H113D) to genetically impair replication-dependent chromatin assembly by destabilizing (H3-H4)_2_ tetramer. We established that DNA synthesis-dependent histone deposition is required for the completion of RDR. Histone deposition prevents joint-molecules from Rqh1-dependent disassembly, a crosstalk contributing to cell survival upon replication stress but channeling RDR towards deleterious events. Asf1 and CAF-1 act in RDR and CAF-1 recruitment to DNA synthesis associated to RDR requires the HR factor Rad52. Our data establish that CAF-1 counteracts Rqh1 activity at sites of replication stress by promoting repair synthesis-coupled histone deposition. These results demonstrate that histone deposition plays an active role in fine-tuning RDR, a benefit counterbalanced by stabilizing at-risk joint molecules for genome stability.

---

- **H3-H113D impairs nucleosome stability and deposition**
- **Histone deposition, Asf1 and CAF-1 play an active role in RDR**
- **RDR-coupled histone deposition impacts genome stability at arrested forks.**
- **CAF-1 is recruited at RDR sites in a Rad52-dependent manner.**

## Introduction

The maintenance of genome integrity occurs in the context of DNA packaged into chromatin. Chromatin constitutes a barrier to DNA replication and repair machineries, that should be first lifted and then restored behind the replication fork or once the repair event is achieved (Soria et al. 2012). Genomes are routinely exposed to a variety of DNA damages that induce profound chromatin rearrangements and pose serious threat to epigenome integrity during DNA replication (Svikovic and Sale 2017). Despite the recent identification of chromatin restoration pathways upon DNA repair, the crosstalk and coordination between both processes, that is likely key to safeguard genome integrity, remain poorly understood (Dabin et al. 2016).

The basic unit of chromatin is the nucleosome which consists of 147 bp of double stranded DNA wrapped around a histone octamer containing one (H3-H4)_2_ tetramer and two H2A-H2B dimers (Luger et al. 1997). During DNA replication, nucleosomes ahead of the replication fork are evicted and both parental and newly synthetized histones are assembled onto newly replicated DNA through a process called replication-coupled chromatin assembly. This process requires a network of chromatin factors that operate sequential reactions to handle histone dynamics at ongoing forks. Nucleosome assembly occurs as a stepwise process in which the (H3-H4)_2_ tetramer is deposited before two H2A-H2B dimers (Smith and Stillman 1991; Hatakeyama et al. 2016). Deposition of (H3-H4)_2_ tetramer requires specific histone modifications and H3-H4 chaperones, such as the Chromatin Assembly Factor 1, CAF-1, the Anti-Silencing Factor 1, Asf1 and Rtt106 (Burgess and Zhang 2013).

CAF-1 plays a key role in nucleosome assembly coupled to DNA synthesis during DNA replication and repair. It associates with the proliferating cell nuclear antigen (PCNA), the processivity factor for DNA polymerases, to facilitate nucleosome deposition onto DNA *in vitro* (Shibahara and Stillman 1999; Moggs et al. 2000). CAF-1 is a tri-subunit complex in which the large subunit (human p150, *S. cerevisiae* Cac1 and *S. pombe* Pcf1), scaffolds interaction with H3-H4 and DNA to allow nucleosome assembly. Recent *in vitro* studies have elucidated how CAF-1 promotes (H3-H4)_2_ tetramer deposition onto DNA (Liu et al. 2016; Zhang et al. 2016; Mattiroli et al. 2017a; Mattiroli et al. 2017b; Sauer et al. 2017). One CAF-1 complex binds a single H3-H4 heterodimer, allowing unmasking the C-terminus winged helix domain of p150 to bind DNA. Then, DNA-mediated dimerization of two CAF-1 complexes allows (H3-H4)_2_ tetramer formation and deposition onto DNA. (H3-H4)_2_ tetramerization is required to achieve deposition onto DNA and then release H3-H4 from CAF-1. An histone H3 mutant that destabilizes H3-H3’ interface impairs *in vitro* tetramer deposition (Sauer et al. 2017). Asf1 binds a H3-H4 heterodimer and acts by transferring H3-H4 to CAF-1 and Rtt106 (English et al. 2006). In yeast models, Asf1 is required for acetylation of H3 at lysine K56 (H3K56Ac), a mark of newly synthetized H3, by the acetyl transferase Rtt109 (Han et al. 2007; Xhemalce et al. 2007). Also, Asf1 associates with components of the replication machinery and facilitates CAF-1-mediated histone deposition *in vitro* (Franco et al. 2005; Burgess and Zhang 2013).

Flaws in the DNA replication process are a source of genome and epigenome instability. Numerous Replication Fork Barriers (RFBs) and replication-blocking agents interrupt fork elongation, causing recurrent temporary pauses to a single replisome and occasional terminal fork arrest. Stressed forks are fragile structures prone to chromosomal aberrations which may result from faulty replication-based DNA repair events (Lambert and Carr 2013). Chromatin establishment and maturation takes place during DNA replication, a critical step to the inheritance of the epigenome (Alabert and Groth 2012). Histone supply and chromatin assembly regulate fork stability and elongation (Mejlvang et al. 2014; Prado and Maya 2017). Fork obstacles interfere with histone dynamics, including histone recycling and inheritance of histone marks, resulting in adjacent loci liable to epigenetic changes (Jasencakova et al. 2010; Svikovic and Sale 2017). Thus, stressed forks are instrumental in triggering chromosomal aberrations and chromatin changes by mechanisms that remain to be fully understood.

A variety of DNA repair factors are engaged in the timely resumption of fork elongation. Homologous recombination (HR) is a key DNA repair pathway that preserves fork integrity and replication competence through a process called Recombination-Dependent Replication (RDR) (Carr and Lambert 2013). At the pre-synaptic step, the recombinase Rad51 is recruited onto single stranded DNA (ssDNA) exposed at arrested forks, to form a filament with the assistance of mediators such as yeast Rad52 and mammalian BRCA2. After homology search at the presynaptic step, the Rad51 filament promotes strand invasion into an intact homologous DNA template, usually the sister chromatid or the parental DNA ahead of the fork, to form a displacement loop (D-loop). The invading 3’ end then primes DNA synthesis to restart the fork and complete DNA replication. D-loops can be disassembled by DNA helicases such as the human RecQ helicase BLM and its fission yeast orthologue Rqh1 (Mimitou and Symington 2009). Because eukaryotic genomes contain several types of dispersed and repeated sequences, RDR can occasionally generate chromosomal rearrangements. In these circumstances, RecQ helicases are instrumental to limit the likelihood of faulty RDR creating chromosomal aberrations (Lambert et al. 2010; Hu et al. 2013).

The access-prime-restore model has put forward the role of chromatin factors to handle histone dynamics at DNA lesions and to initiate DNA repair, with mammalian Asf1 and CAF-1 being involved in the early step of DSB repair by HR (Soria et al. 2012). Subsequent to DNA repair, chromatin restoration is a necessary step to engage physiological processes such as transcription restart and turning-off checkpoint response (Chen et al. 2008; Adam et al. 2013). Crosstalk to couple DNA repair and chromatin re-assembly are poorly understood and it is unknown whether RDR remains coupled to histone deposition in S-phase. We previously reported that fission yeast CAF-1 is required to complete the process of RDR in a PCNA-dependent manner (Pietrobon et al. 2014). By counteracting Rqh1-dependent disassembly of D-loops, CAF-1 channels RDR events towards a chromosomal rearrangement pathway. However, the mechanism by which CAF-1 counteracts Rqh1 activity at site of replication stress is unknown.

Using a well-established and characterized RDR assay in fission yeast, we have found that RDR is coupled to Asf1 and CAF-1-mediated histone deposition, a step necessary to RDR completion. We established that RDR-coupled histone deposition prevents the disassembly of D-loops by Rqh1, a crosstalk promoting cell resistance upon replication stress but channeling HR events towards deleterious recombinants types. Moreover, CAF-1 is recruited at sites of DNA synthesis associated to RDR in a Rad52-dependent manner, indicating that CAF-1-mediated histone deposition takes place downstream from the initial step of RDR. Therefore, we demonstrate that CAF-1 counteracts Rqh1 activity by promoting histone deposition during the DNA repair synthesis step associated to RDR. Altogether, our data reveal a novel crosstalk between DNA repair factors and chromatin re-assembly which balances genome stability at site of fork-arrest.

## Results

### Asf1 is required to complete RDR

We previously reported that CAF-1 acts during the DNA synthesis step of RDR, in a way that D-loops are protected from disassembly by Rqh1. Here, we asked whether Asf1 is also involved in RDR. To this end, we employed a previously described site-specific fork arrest assay in which replication of a specific genomic locus is dependent on HR (Lambert et al. 2010). The assay consists of a polar Replication Fork Barrier (RFB), called *RTS1*, integrated as inverted repeats at both sides of the *ura4^+^* gene, abbreviated as the *t>ura4<ori* locus (Figure 1A). Upon activation of the RFB, the binding of Rad51 and its Rad52 loader allows blocked forks to be restarted to overcome the RFB. Occasionally, Rad51 promotes newly replicated strands to switch template and invades the opposite *RTS1* sequence to form a D-loop which primes the restarted DNA synthesis on a non-contiguous DNA template. This faulty restart pathway results in the formation of stable joint-molecules (JMs), referred to as D-loop and Holliday junction-like intermediates, whose resolution generates at least two RDR products: an acentric and a dicentric chromosome (Pietrobon et al. 2014). The formation of acentric and dicentric is strictly dependent on Rad52 and serves as a marker for RDR completion (Lambert et al. 2010).

**Figure 1:**
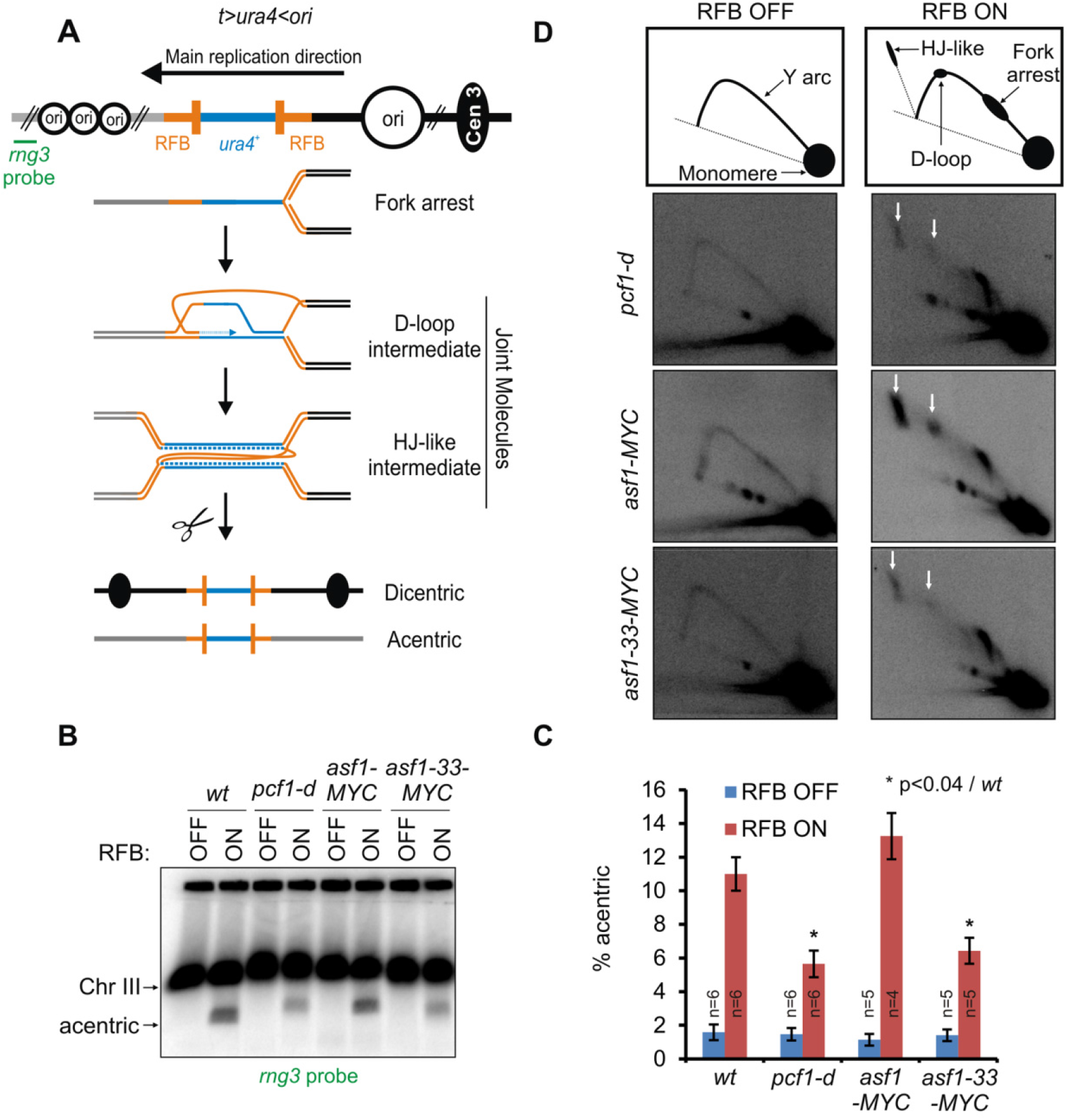
Loss of Asf1-function impairs RDR. **A**. Diagram of the *t>ura4<ori* locus, in which *t* refers to the telomere-proximal side (gray lines), *ura4* refers to the *wt* gene (blue line), > *and <* refers to the polarity of the two *RTS1-RFBs* (orange bars) and *ori* refers to the replication origin (opened black circle, the largest one being the most efficient origin). Green bar indicates the *rng3* probe. The RDR assay consists of a polar Replication Fork Barrier (RFB), called *RTS1*, integrated at the *ura4^+^* gene, 5Kb away from a strong replication origin at the centromere-proximal side. An inverted repeated *RTS1* sequence is integrated at the telomere-proximal side of *ura4^+^* to generate the *t>ura4<ori* locus. Due to the main replication direction, the barrier activity of the centromere-proximal RTS1-RFB is predominant over the activity of the telomere-proximal RFB. The RFB activity is mediated by the RTS1-bound protein Rtf1, the expression of which is regulated by the *nmt41* promoter repressed in the presence of thiamine. Upon Rtf1 expression, > 90% of forks are blocked at the centromere-proximal RFB. The binding of the Rad51 recombinase and its Rad52 loader allow the blocked fork to be restarted to overcome the RFB. Faulty restart events occur in ~ 2-3 % of cells/replication: Rad51 promotes newly replicated strands to switch template and to invade the opposite inverted *RTS1* sequence. DNA synthesis is then initiated from the 3’ invading strand on a non-contiguous DNA template, resulting in a stable early JM, referred to as a D-loop intermediate. Upon arrival of the converging fork, a second template switch event results in the formation of a later JM, referred to as Holliday junction (HJ)-like intermediate whose resolution generates at least two recombination products: acentric and dicentric which levels are a marker of RDR completion. **B**. Chromosome analysis in indicated strains and conditions by PFGE and Southern-blot using a radiolabeled *rng3* probe. C. Quantification of acentric level normalized to chromosome III level. Values are means of n independent biological replicates ± standard error of the mean (SEM). Statistical analysis was performed using Mann & Whitney U test. D. Top panel: schematics of replication intermediates (RIs) observed by 2DGE, in RFB OFF and ON conditions. Bottom panels: representative 2DGE experiments in indicated strains and conditions. White arrows indicate JMs. See Figure S1 for RDR analysis in additional chromatin factor mutants.

We applied the RDR assay to a thermo-sensitive allele of the essential *asf1* gene, *asf1-33*, which was reported to be defective for H3 deposition at restrictive temperature (36°C, (Tanae et al. 2012)). Cells were cultured at semi-permissive temperature (32°C) at which *asf1-33* mutated cells exhibit sensitivity to methyl methane sulfonate (MMS, an alkylated DNA agent that impedes replication fork progression) and a reduced Asf1-H3 interaction (Figure S1A-B). Chromosome analysis by Pulsed Field Gel Electrophoresis (PFGE) coupled with Southern-blot hybridization showed that the amount of acentric fragment, a RDR product, was reduced in *asf1-33* cells, as well as in a mutant lacking CAF-1 (*i.e*. in *pcf1* deleted cells), indicating that Asf1 promotes RDR (Figure 1B-C). The analysis of replication intermediates by two-dimensional gel electrophoresis (2DGE) showed that signals corresponding to arrested forks were similar in all analyzed strains, indicating that defect in RDR is not a consequence of a less efficient RFB activity in strains lacking CAF-1 or Asf1. We observed that JMs intensity was similarly reduced in *asf1-33* and *pcf1-d* mutants (Figure 1D), showing that Asf1 preserves JMs during RDR, as proposed for CAF-1. It is worth noting that the *asf1-33* mutated cells form Rad52 foci, indicating that Asf1 is dispensable to the early step of HR (Tanae et al. 2012). As previously proposed for the lack of CAF-1 (Pietrobon et al. 2014), these data suggest that JMs are faster disassembled upon loss of Asf1 function rather than being less efficiently formed. We concluded that Asf1 and CAF-1 are necessary to RDR completion, possibly by promoting histone deposition during the DNA synthesis step of RDR, thus creating a substrate less favorable to Rqh1-dependent D-loop disassembly.

We investigated the role of other histone chaperones in RDR by analyzing the level of acentric chromosome upon RFB induction. We found no impact on acentric level in cells lacking the HIRA complex (involved in replication-independent H3-H4 deposition), the FACT complex, Rtt106, the two orthologues of the *S. cerevisiae* H2A-H2B histone chaperone Nap1 and Nap2, and Chz1, a histone variant H2AZ chaperone (Figure S1C-D). We concluded that RDR requires the two key histone chaperones Asf1 and CAF-1 which mediates histone deposition in a DNA-synthesis dependent manner.

### The mutated histone H3-H113D disrupts (H3-H4)_2_ tetramer formation

Our data suggest a scenario in which Asf1 and CAF-1 promote chromatin assembly during the DNA synthesis step of RDR to prevent D-loop disassembly. To address the role of histone deposition during RDR, we decided to genetically disrupt replication-dependent chromatin assembly by altering the stability of (H3-H4)_2_ tetramer to inhibit their stable deposition onto DNA. To this end, we employed a mutated form of H3 containing a single substitution of histidine to aspartic acid (H3-H113D). This mutation was reported to impair CAF-1-mediated nucleosome deposition *in vitro* (Nakano et al. 2011).

The interface between the two H3-H4 dimers involves the C-terminal region (residues 106 to 131) of the two histones H3, called here H3 and H3’ (Luger et al. 1997; Davey et al. 2002) (Figure 2A-B and S2A). We examined X-ray structures of nucleosomes, chosen among the numerous available experimental models on the basis of homology with *S. pombe* histone H3 (see details in materials and methods). The H113 emerges as a key residue in the H3:H3’ interface with H113 of one histone H3 being anchored to the second histone H3’ by a dense network of contacts involving six residues (Figure S2B). Each H113 forms two intermolecular hydrogen bonds with C110 and D123, reinforced by Van der Waals contacts with four residues: A114, R116, K122 and L126. By replacing a neutral or positively charged histidine residue by a negatively charged aspartate that is positioned in front to another aspartate, D123, the H113D mutation generates a prohibitive electrostatic repulsion in the intact H3:H3’ organization. Previous works reported that a series of mutation (C110E, L12R-I130R, H113A, L126A and A114Y) prevent the H3:H3’ interface formation because of an overwhelming energetic penalty and such mutations are lethal in budding yeast (Banks and Gloss 2004; Ramachandran et al. 2011; Sauer et al. 2017). By analogy with these cases, the H113D mutation likely drastically destabilizes the H3:H3’ interface, precluding the (H3-H4)_2_ tetramer formation.

**Figure 2:**
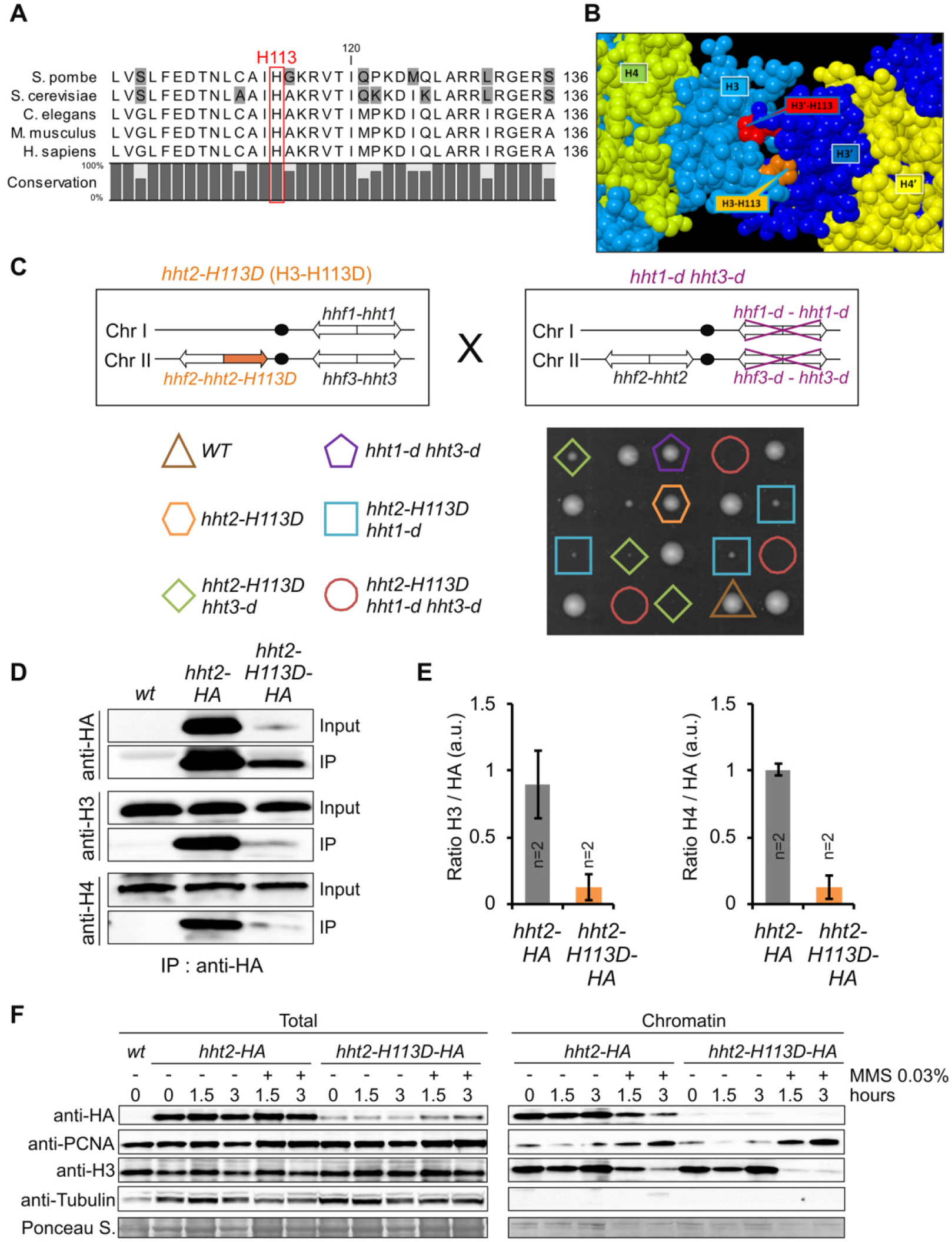
The mutated histone H3-H113D destabilizes (H3-H4)_2_ tetramer. **A**. Alignment of yeast, worm, mouse and human histone H3. Red rectangle indicates the position of histidine 113. UniProtKB access codes used are *S.pombe*: P09988; *S. cerevisiae*: P61830; *C.elegans*: P08898; *M. musculus*: P84228 and *H. sapiens*: Q71DI3. **B**. View of the H3:H3’ interface. The interface between the two histones H3, colored here in light (H3) and dark blue (H3’), maintains H3 and H4 in the tetramer form (H3-H4)_2_. In the H3:H3’ interface, H113 of each histone (orange for H113 in H3 and red for H113 in H3’) is deeply buried in the adjacent histone partner. For clarity, the other histones H2A and H2B, as well as DNA, are not represented. **C**. Spore viability analysis of indicated genotypes. **D**. Association of H3-HA and H3-H113D-HA with untagged H3 and H4 in indicated strains. **E**. Quantification of panel D expressed in arbitrary unit (a.u.). Values are means of n independent biological replicates ± standard deviation (SD). **F**. Chromatin association of analyzed proteins in indicated strains and conditions (hours upon MMS addition or not). See Figure S2 for structural impact of H3-H113D.

In *S. pombe* cells, 3 genes *(hht1, hht2* and *hht3)* encode a single histone H3 protein. The H113D mutation was introduced in the *hht2* gene and the resulting H3-H113D expressing cells were viable with no apparent growth defect (Figure 2C). The deletion of *hht1* and *hht3* is viable *(hht1-d hht3-d)* and cells maintain histone H3 protein levels similar to those in *wt* strain (Yadav et al. 2017). To obtain a strain expressing only H3-H113D, the *hht2-H113D* mutated strain was crossed with an *hht1-d hht3-d* strain. We found that combining the *hht2-H113D* mutation with both deletions resulted in a synthetic lethality (Figure 2C). Thus, when H3-H113D is the sole cellular histone H3 expressed, cells are not viable, in agreement with the hypothesis that this mutation inhibits (H3-H4)2 tetramer formation. Still, we observed that combining *hht2-H113D* with either single *htt1* or *hht3* deletion preserves cell viability but causes a severe synthetic growth defect (Figure 2C). This strongly suggests that *wt* H3 is abundant enough to allow *wt* (H3-H4)_2_ tetramer to be formed and to preserve cell viability.

We probed H3-H113D associations with histones H3 and H4 in *hht2-H113D* mutated cells *(hht1^+^ hht3^+^)*. Although, the protein level of H3-H113D-HA (an HA tagged version of the mutated histone) was lower than the one of *wt* H3-HA (Figure 2D and S2C), immuno-precipitation experiments from total protein extracts clearly showed that H3-H113D-HA association with *wt* H3 and H4 were severely reduced, compared to H3-HA (Figure 2D-E). These data indicate that mixed (H3-H113D-H4-H3-H4) tetramers are highly unstable. We asked if H3-H113D is incorporated into chromatin using chromatin fraction assay. As expected, the *wt* H3-HA was found to be chromatin-bound whereas H3-H113D-HA was not detected in the chromatin fraction, indicating that this mutated histone is very poorly incorporated into nucleosomes assembled onto DNA (Figure 2F). *Wt* H3 (expressed from *hht1* and *hht3* gene) was equally detected in the chromatin fraction of *hht2-HA* and *hht2-H113D-HA* cells, indicating that *wt* H3 is *in fine* incorporated within *wt* nucleosomes in H3-H113D expressing cells. Altogether, our data indicate that in *hht2-H113D* mutated cells, H3-H113D inhibits stable tetramer formation and is therefore not deposited onto chromatin, while *wt* (H3-H4)2 tetramers are formed and assembled onto chromatin.

### H3-H113D impairs replication-coupled nucleosome assembly

H3-H113D inhibits tetramers formation and is not deposited onto chromatin. We thus investigated its consequences on replication-coupled chromatin assembly pathways. Asf1-Myc associated with *wt* H3 and H3-HA but not with H3-H113D-HA (Figure 3A and S3A), arguing that the histone chaperone function of Asf1 is preserved in *hht2-H113D* mutated cells. We found that H3-H113D-HA binds to Pcf1-YFP, without preventing Pcf1-YFP to associate with Pcf2-MYC (the second CAF-1 subunit), PCNA and *wt* H3; these protein-protein interactions being necessary for optimal CAF-1-mediated histone deposition (Figure 3A and S3B-C). We concluded that in *hht2-H113D* mutated cells, CAF-1 forms complexes with H3, H3-H113D, and PCNA. Because H3-H113D inhibits tetramer formation and that tetramerization is necessary to CAF-1 mediated histone deposition *in vitro* (Sauer et al. 2017), H3-H113D may impair CAF-1-dependent chromatin assembly *in vivo*.

**Figure 3:**
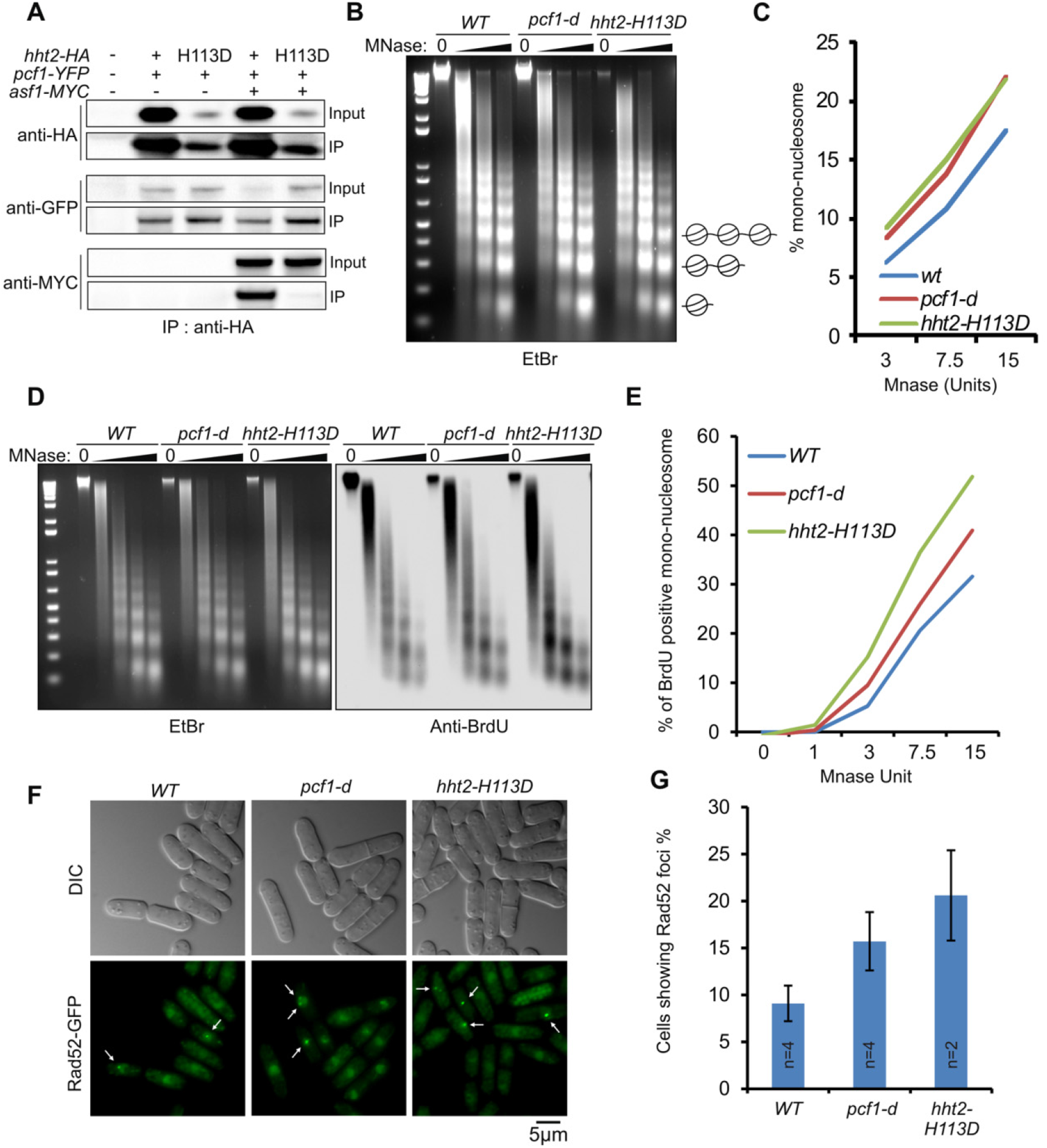
H3-H113D impairs replication-coupled chromatin assembly. **A**. Association of H3-HA and H3-H113D-HA with Pcf1-YFP and Asf1-MYC in indicated strains. **B**. Representative experiment of chromatin digestion by MNase. Genomic DNA of wild type and mutant strains was digested with increasing amount of MNase and migrated on ethidium bromide (EtBr)-containing agarose gel. **C**. Percentage of mono-nucleosome relative to total DNA. **D**. Representative experiment of replicated chromatin sensitivity to MNase treatment in indicated strains. Left panel: BrdU-incorporated genomic DNA was digested with increasing amount of MNase and migrated on ethidium bromide-containing agarose gel. Right panel: after transfer onto nitrocellulose membrane, incorporated BrdU was revealed using anti-BrdU antibody. **E**. Percentage of BrdU-positive mono-nucleosome relative to total BrdU signal. **F**. Representative examples of Rad52-GFP foci (white arrows) in indicated strains. G. Quantification of E. Values are means of n independent biological replicates ±SEM. See Figure S3 for protein-protein interactions and S4 for biological replicates of sensitivity to MNase.

We further investigated the impact of H3-H113D on chromatin structure. We analyzed the MNase sensitivity of chromatin from *wt, pcf1-d* and *hht2-H113D* cells. Compared to *wt* strain, the proportion of mono-nucleosome population was increased in *pcf1-d* and *hht2-H113D* cells, showing a higher sensitivity to MNase as an indication of decreased global nucleosomal density (Figure 3B-C and S4A-B). To test if this chromatin changes are caused by defective replication-coupled chromatin assembly, we analyzed the MNase sensitivity of the replicated chromatin. The *hht2-H113D* mutation was introduced in strains able to incorporate BrdU, a thymidine analogue (Fleck et al. 2017). Cells were blocked in early S-phase by hydroxyurea treatment and released in BrdU-containing media for 20 minutes to label the replicated chromatin. BrdU incorporation was detected in MNase-digested chromatin only when cells were released from HU block, showing that BrdU-labelling marks the replicated chromatin (Figure S4C). Compared to *wt* strain, the proportion of BrdU-positive mono-nucleosome was increased in *hht2-H113D* and *pcf1-d* mutated cells, indicating that the chromatin associated to newly replicated strands is more sensitive to MNase digestion (Figure 3D-E and S4D-E). These data are consistent with a defect in replication-coupled chromatin assembly in the absence of CAF-1 and in H3-H113D expressing cells. Previous reports in budding yeast showed that defective histone deposition behind replication forks results in DSBs (Clemente-Ruiz and Prado 2009). Consistent with this, *pcf1-d* and *hht2-H113D* mutated cells exhibited a high frequency of cells showing Rad52 foci, a marker of DNA lesion (Figure 3F-G). Of note, H3-H113D expressing cells are competent to form Rad52 foci, indicating that the presynaptic steps of the HR process are likely unaffected when inhibiting (H3-H4)_2_ tetramer formation and deposition. We concluded that H3-H113D mutation impairs replication-coupled histone deposition to an extent at least similar to a CAF-1 defect. Although other histone deposition pathways might be affected, our data are consistent with the fact that in H3-H113D expressing cells, the CAF-1-mediated chromatin assembly pathway is impaired.

### Histone deposition acts in RDR to prevents D-loop disassembly

The H3-H113D offers the possibility to question the role of replication-coupled chromatin assembly during RDR. We thus applied the RDR assay to *hht2-H113D* mutated cells and found that JMs intensity was reduced as well as the subsequent accumulation of acentric chromosome, one RDR product (Figure 4A-C). These data indicate that histone deposition is required to RDR completion by preserving JMs. No additive effect was observed in the *pcf1-d hht2-H113D* strain (Figure 4A-B), showing that CAF-1 and histone deposition act in a same pathway of promoting RDR. We reported that in the absence of CAF-1, JMs are faster disassembled by Rqh1 (Pietrobon et al. 2014). Remarkably, similar interactions were observed in *hht2-H113D* cells in which the deletion of *rqh1* restored the intensity of JMs signal (Figure 4C). Since the presynaptic steps of the HR process appear functional in *hht2-H113D* cells (Figure 3F-G), these data strongly support that inhibiting (H3-H4)_2_ tetramer assembly onto DNA favors D-loop disassembly by Rqh1. Consistently, no defect in RDR were observed in strains in which genes encoded H3 are deleted (either *hht2-d* or *hht1-d hht3-d* cells, Figure 4A and S5), indicating that H3-H113D favors D-loop disassembly as a consequence of impairing replication-coupled histone deposition. Altogether, these data establish that histone deposition coupled to RDR is required to prevent the disassembly of the D-loop by Rqh1.

**Figure 4:**
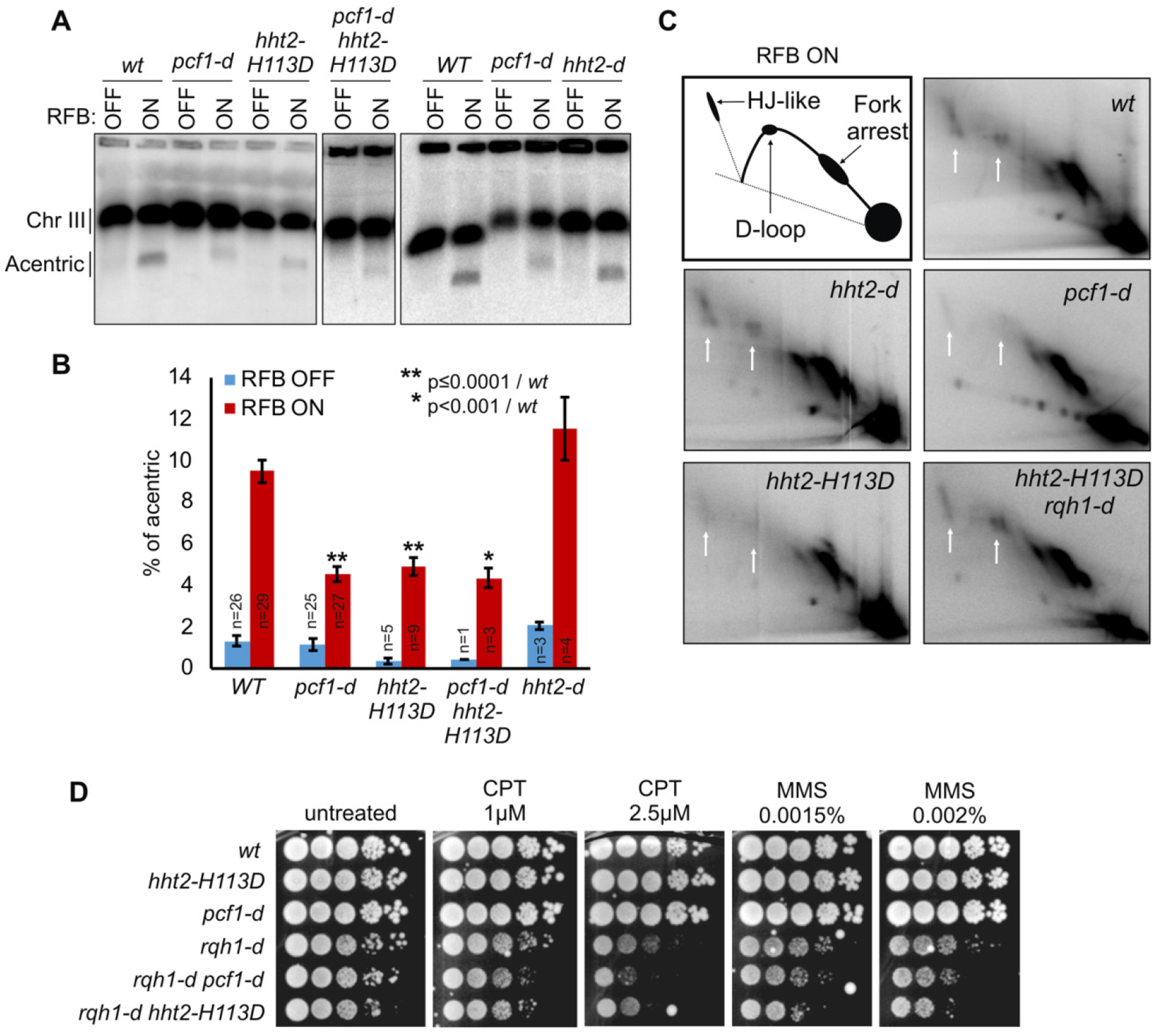
Histone deposition coupled to RDR preserves JMs. **A**. Chromosome analysis in indicated strains and conditions as described on Figure 1. **B**. Quantification of acentric level as described on Figure 1. Values are means of n independent biological replicates ± SEM. Statistical analysis was performed using Mann & Whitney U test. **C**. Representative 2DGE analysis in indicated strains. White arrows indicate JMs. **D**. Survival assay in 10fold serial dilution experiment of wild type and mutant strains in indicated conditions. See Figure S5 for the role of H3K56Ac in RDR.

### Crosstalk between Rqh1 and RDR-coupled histone deposition promotes resistance to replication stress

We reported that genetic interactions between CAF-1 and Rqh1 promote survival to various replication-blocking agents. We observed that H3-H113D mutation increased the cell sensitivity of *rqh1-d* deleted cells to MMS and campthotecin (CPT, a topoisomerase I inhibitor), thus mimicking CAF-1 defect (Figure 4D). This was not a consequence of a loss of Rqh1-Pcf1 interaction as Rqh1-MYC associated with Pcf1-YFP in *hht2-H113D* mutated cells (Figure S3C). Also, MMS treatment did not stimulate chromatin-bound H3-H113D-HA, suggesting that unstable tetramers are unlikely to be assembled during repair synthesis (Figure 2F). Altogether, our data indicate that the antagonistic activities of CAF-1-mediated histone deposition and Rqh1 in JMs resolution during RDR contribute to cell resistance to replication stress.

### RDR-coupled histone deposition favors deletion type recombinant

Our data reveal antagonistic roles of RDR-coupled histone deposition and Rqh1 in JMs resolution. We investigated the consequences of this crosstalk on HR outcomes, by monitoring spontaneous HR events using an assay for intra-allelic recombination between direct *ade6* (Figure 5A) (Hartsuiker et al. 2001). Gene conversion (GC) and synthesis dependent strand annealing (SDSA) result in conversion type, whereas crossover (CO) and single strand annealing (SSA, a DNA repair pathway independent of Rad51-mediated strand exchange) give rise to deletion type. In *rqh1-d* cells, conversion and deletion events were increased by 2- and 3-fold, respectively, which is consistent with known Rqh1 anti-recombinase activity (Figure 5B) (Doe et al. 2000). The H3-H113D mutation had little impact on HR outcomes but when combined with *rqh1-d*, it partially rescued the deletion rate, and suppressed bias towards deletion type. These data are consistent with antagonistic activities of Rqh1 and histone deposition in controlling HR outcomes: Rqh1 promotes D-loop disassembly to channel HR event towards a conservative pathway whereas histone deposition favors D-loop stability to channel HR event towards deletion events such as CO.

**Figure 5:**
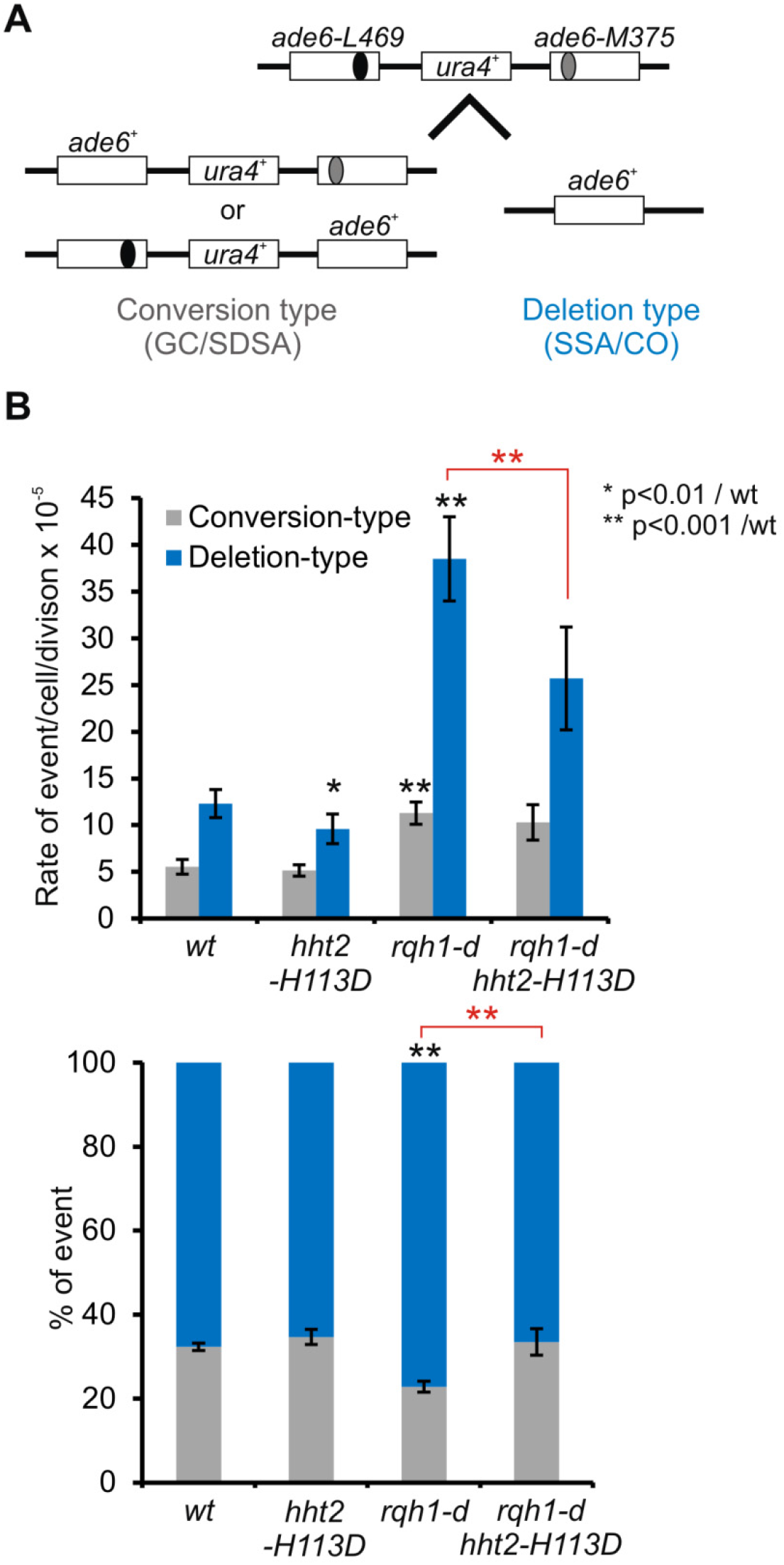
Histone deposition channels HR towards deletion-type events. **A**. Schematic representation of HR substrate and recombination outcomes. **B**. Top panel: rate of conversion and deletion type in indicated strains. Values are median rate calculated from at least 25 independent cultures ± 95 % confidence interval (CI). Bottom panel: ratio of deletion and conversion type in indicated strains. Error bars indicate SEM. Statistical analysis was performed using Mann & Whitney U test. Black and red stars indicate statistics compared to *wt* and compared to *rqh1-d*, respectively.

### H3K56Ac is dispensable to RDR

The H3K56Ac modification marks newly synthetized histone and was proposed to regulate nucleosome assembly and to contribute to the DNA damage response (Driscoll et al. 2007). Hence the importance of acetylation in addition to histone deposition has to be considered. We found that in contrast to *wt* H3 which was efficiently acetylated by Rtt109 in *hht2-H113D* cells, H3-H113D that is unable to associate with Asf1-MYC (Figure 3A) lost the H3K56Ac mark (Figure S5A). We thus tested the role of H3K56Ac in RDR. We analyzed two strains: one expressing a single H3 protein which cannot be acetylated on K56 *(hht1-d hht3-d hht2-K56R)* and a *rtt109* deleted strain. Consistent with previous reports (Xhemalce et al. 2007), both strains were defective for H3K56Ac (Figure S5B) and were proficient for the formation of acentric chromosome, a RDR product (Figure S5C-D). Thus, H3K56Ac is dispensable to RDR completion.

### CAF-1 recruitment to site of DNA synthesis during RDR requires Rad52

Our data are consistent with RDR being coupled to histone deposition mediated by the Asf1 and CAF-1 chaperones. We investigated the recruitment of CAF-1 and histone H3 to damaged chromatin using chromatin fraction assays. The chromatin-bound Pcf1 fraction was enriched after treatment with MMS, but not with bleomycin, a DSB-inducing agent, or hydroxyurea (an agent that slows down fork progression) (Figure 6A). Consistent with CAF-1 being a scarce complex (~ 500 to 900 molecules/cells (Carpy et al. 2014), Pcf1-YFP and Pcf2-MYC were hardly detectable in the total fraction, but were clearly enriched in the chromatin fraction from 1-2 hours after MMS treatment (Figure 6B-C). This recruitment was concomitant with PCNA, the factor that links CAF-1 function to the step of DNA synthesis during RDR (Pietrobon et al. 2014). It is worth noting that in *rad52-d* mutant, upon MMS treatment, the chromatin-bound PCNA and H3 fractions were reduced (Figure S6A), indicating that the HR process is necessary for PCNA recruitment and the retention/restoration of histone on damaged chromatin. However, neither H3-H4 nor PCNA enrichment to damaged chromatin were affected by the lack of CAF-1 (Figure S6B), revealing that multiple histone chaperones likely contribute to chromatin restoration in response to MMS.

**Figure 6:**
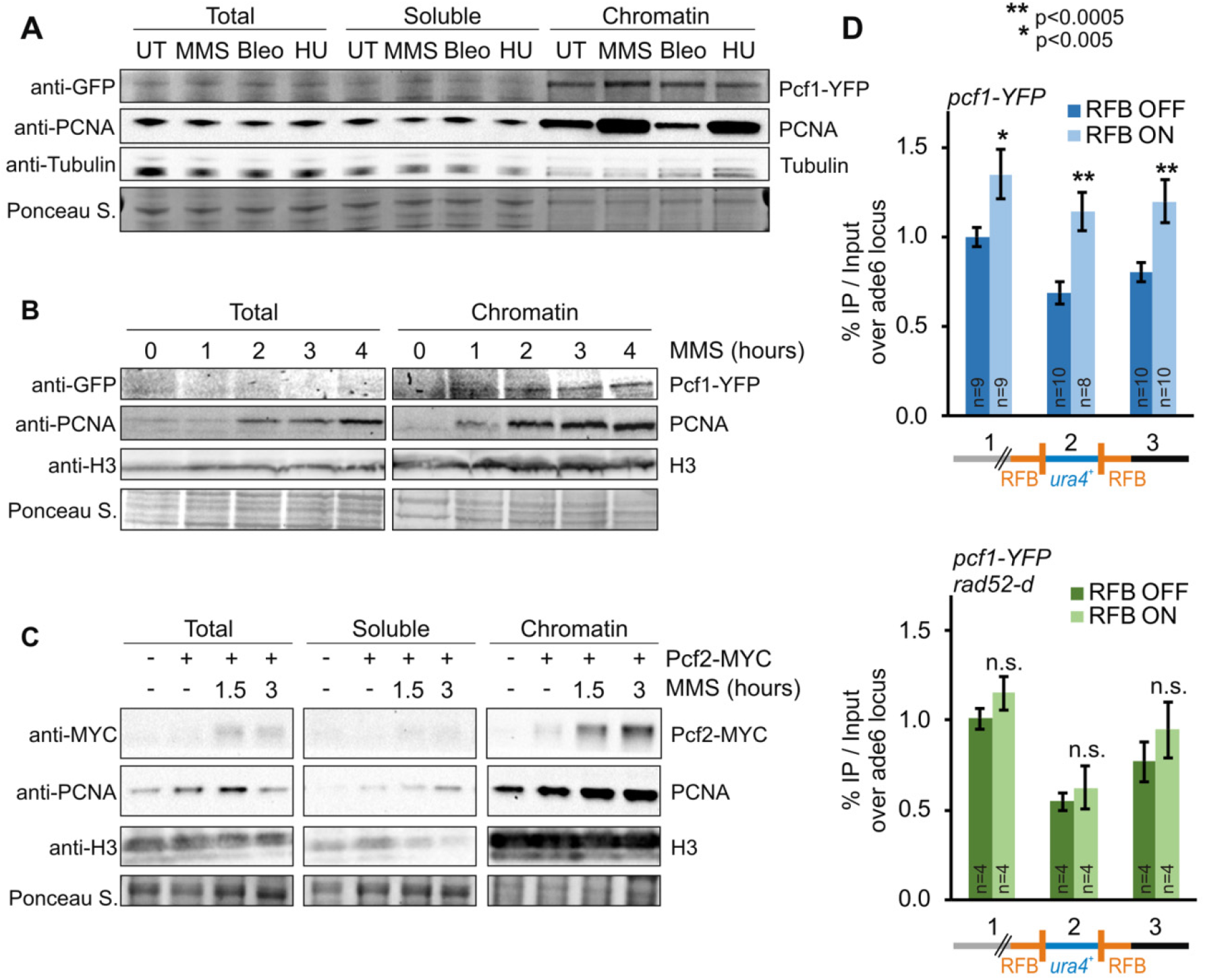
CAF-1 recruitement to RDR sites requires Rad52. **A**. Chromatin association of Pcf1-YFP in indicated conditions (MMS: 2 hours of 0.03 % MMS, Bleo: 2 hours of 3.5mU/ml of Bleomycin, HU: 2 hours of 20 mM of HU). These doses result in significant cell death in chronic treatment and were thus used in acute treatment. B. Kinetics of chromatin association of Pcf1-YFP after 0.03% MMS treatment. **C**. Kinetics of chromatin association of Pcf2-MYC after 0.03% MMS treatment. **D**. Analysis of Pcf1-YFP binding to the t>ura4<ori locus in RFB OFF and ON conditions (top panel: *wt strain*, bottom panel: *rad52-d strain)*. Scheme at the bottom depicts the primers location within the *t>ura4<ori* locus: primer pair 1 are located 400 bp at the telomere-proximal side, primer pair 2 are within the *ura4* gene, primer pair 3 are located 150 bp at the centromere-proximal side. Values are means of n independent biological replicates from 2 independent clones ± 95 % CI. Statistical analysis was performed using Mann & Whitney U test. See Figure S6 for PCNA and H3 association to chromatin in *rad52-d* mutant.

Due to technical issues, we were unable to generate chromatin fractions of good quality in *rad52-d* cells, which precluded us from addressing whether or not HR factors are required for CAF-1 recruitment to damaged chromatin. Thus, we took advantage of the RDR assay and found that Pcf1-YFP was recruited at the two active RFBs (Figure 6D, primer pairs 1 and 3, top panel). Upon activation of the RFBs, the DNA synthesis of the *ura4* gene occurs through RDR. We observed that Pcf1-YFP was recruited to *ura4* only in “ON” condition (primer pair 2, top panel), showing that Pcf1 binds to sites of DNA synthesis occurring during RDR. This Pcf1 binding was no longer observed in *rad52-d* cells in which RDR and JMs cannot occur (Figure 6D, bottom panel) (Lambert et al. 2010). We concluded that CAF-1 recruitment to RDR sites occur downstream from the early step of RDR and JMs formation.

## Discussion

Fork-obstacles open up the risk of genome and epigenome instability. We have established that RDR, a major fork-restart pathway, is coupled to histone deposition, mediated by the chromatin assembly factors CAF-1 and Asf1 (Figure 7). We made the surprising finding that histone deposition coupled to restarted DNA synthesis impacts the subsequent resolution of JMs at site of fork arrest, and as a consequence favors the channeling of RDR events towards a chromosomal rearrangement pathway. We reveal a novel crosstalk between chromatin restoration and a DNA repair pathway which is required to balance genome stability at arrested forks and to cell survival upon replication stress. Also, our finding highlight that chromatin restoration is entirely part of the RDR process. We proposed a scenario in which histone deposition plays an active role in RDR to avoid discontinuity in chromatin assembly upon replication stress, a benefit counterbalanced by stabilizing at-risk joint molecules for genome stability.

**Figure 7:**
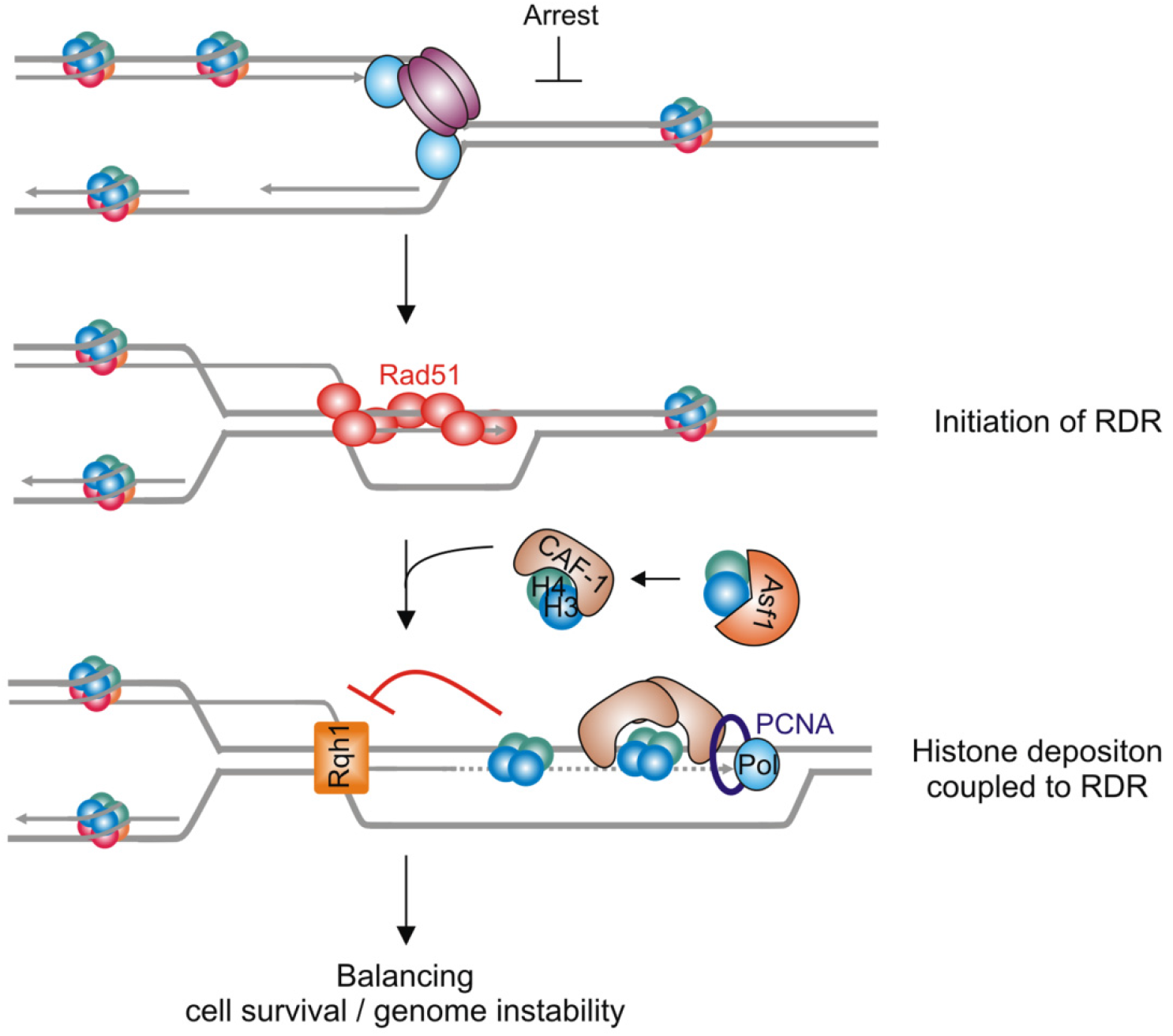
Model of histone deposition coupled to recombination-dependent replication. Upon fork arrest, HR factors promote D-loop formation which primes the restarted synthesis and histone deposition. Histone deposition coupled to RDR allows JMs to be protected from disassembly by Rqh1. This antagonistic activity of histone deposition and Rqh1 in JMs protection/disassembly contributes to cell resistance to replication stress and balance genome stability at site of replication stress.

### Asf1, CAF-1 and histone deposition act in RDR at the step of DNA synthesis

We previously reported that CAF-1 acts in RDR in a way that D-loops are less efficiently disassembled by Rqh1 (Pietrobon et al. 2014). Despite recent advances in understating the mechanism of CAF-1-mediated histone deposition *in vitro*, it remains challenging to define point mutations to generate mutated forms of CAF-1 unable to interact with H3-H4 *in vivo* (Sauer et al. 2017). To overcome this, we took advantage of the H3-H113D mutated form, reported to inhibit CAF-1-mediated histone deposition *in vitro* (Nakano et al. 2011). We show here that the H3-H113D precludes (H3-H4)_2_ tetramer formation, and is thus poorly incorporated into nucleosomes deposited onto DNA and inhibits replication-coupled chromatin assembly. Remarkably, the H3-H113D mutation mimics the absence of CAF-1 in impairing RDR. RDR also requires Asf1, the three CAF-1 subunits and the ability of CAF-1 to interact with PCNA (Pietrobon et al. 2014). Altogether, these data support the view that Asf1 and CAF-1 fine-tune RDR by coupling histone deposition to the step of DNA synthesis. Consistent with this, CAF-1 binds to sites of HR-dependent DNA synthesis in a Rad52-dependent manner, indicating that the early steps of RDR must be engaged for the subsequent recruitment of CAF-1.

Mammalian CAF-1 and ASF1 are necessary for the early steps of recombinational repair of DSBs by facilitating the loading of HR factors (Baldeyron et al. 2011; Huang et al. 2018). Our data favors a model in which fission yeast CAF-1 and histone deposition act in the later steps of RDR. Firstly, Rad52 was able to bind arrested forks in the absence of CAF-1 (Pietrobon et al. 2014). Secondly, the binding of CAF-1 to the sites of DNA synthesis occurs downstream from Rad52. Thirdly, RDR defects are rescued by deleting *rqh1*. This indicates that, in cells lacking the pathway of histone deposition coupled to RDR, D-loops are faster disassembled rather than unable to form. We proposed that CAF-1-mediated histone deposition acts as a chromatin restoration pathway during the DNA synthesis step of RDR. In this scenario, histone deposition would occur onto the DNA duplex of the extended D-loop (Figure 7). However, we cannot exclude that histone deposition occurs onto the displaced strand of the D-loop, as it has been proposed that nucleosome can be deposited onto ssDNA (Adkins et al. 2017). RDR requires two histones chaperones that mediate chromatin assembly in a DNA synthesis manner and requires the ability of CAF-1 to interact with PCNA. Therefore, we favor the first hypothesis in which histone deposition is coupled to DNA synthesis to extend the D-loop thus creating a substrate less favorable to Rqh1-dependent D-loop disassembly. Extensive works have address the role of chromatin factors in regulating genome accessibility to DNA repair machineries but how chromatin restoration is coupled to the DNA repair event is poorly understood (Dabin et al. 2016). Our data put forward a crosstalk between the DNA repair machinery and the step of chromatin restoration. Specifically, we demonstrate that chromatin restoration plays an active role in fine-tuning RDR and in the subsequent resolution of JMs to impact genome stability.

### Histone deposition coupled to RDR impacts genome stability at site of replication stress

Our data establish that CAF-1 counteracts Rqh1 activity at sites of replication stress by promoting repair synthesis-coupled histone deposition. We propose that histone deposition onto extended D-loops creates a substrate less favorable to the Rqh1 activity. RecQ helicases act as motor proteins/helicases to migrate DNA junctions which may be prevented by assembled nucleosomes. It remains challenging to address whether histone are deposited onto D-loops as these structures are very transient. JMs visualization by 2DGE requires a step of enrichment in replication intermediates, technically incompatible with chromatin immuno-precipitation approaches to address histone binding to JMs. Nonetheless, we established that inhibiting replication-coupled histone deposition favors D-loops disassembly. We provide insights into the nature of the chromatin required to protect D-loops (Figure 7).

Asf1 acts as histone chaperone to present H3-H4 to Rtt109 and establish the H3K56Ac mark and then transfer H3-H4 to CAF-1 (Burgess and Zhang 2013). We found H3K56Ac dispensable to promote RDR. Possibly, Asf1 is required for RDR by acting as a donor histone to CAF-1. However, H3-H113D binds to CAF-1, likely as a H3-H4 dimer, but not to Asf1, indicating that, during RDR-coupled histone deposition, H3-H4 can be handed off to CAF-1 independently of Asf1, a pathway that remains to be clarified.

Nucleosome deposition onto extended D-loops may impose topological constraints. D-loops disassembly is a topoisomerase-mediated mechanism (Fasching et al. 2015). Thus, topological constraints resulted from DNA wrapped around nucleosome may be easily relieved by topoisomerase 3. When seeking for additional chromatin factors required for RDR, we found Nap1 and Nap2, two histone H2A-H2B chaperones, to be dispensable to protect D-loops. This suggests that the deposition of (H3-H4)_2_ tetramer, but not the formation of a nucleosome, is sufficient by itself to counteract Rqh1 activity and to limit topological constraints.

During HR-mediated DSB repair, disassembly of D-loops extended by DNA polymerase ensures a noncrossover outcome (Mimitou and Symington 2009). Our data indicate that histone deposition during RDR stabilizes JMs and favors deletion events and a chromosomal rearrangement pathway. Similarly, in the absence of Rqh1, spontaneous HR events are channeled towards deletion type recombinants in a histone deposition manner. Interestingly, spontaneous rates of gene conversion were unaffected by the defect in repair synthesis-histone deposition. This suggests that DNA synthesis associated to GC and SDSA is too short in length to favor histone deposition. Nonetheless, our data reveal that the antagonistic activities of RDR-coupled histone deposition and Rqh1 in D-loops resolution are pivotal to balance genome stability at arrested forks and to promote cell resistance upon replication stress.

### RDR is coupled to chromatin assembly

During unchallenged replication, the concerted action of multiple histone chaperones coordinates the assembly of chromatin behind the fork to achieve recycling of parental histones and deposition of newly synthetized histones (Alabert and Groth 2012). RDR results in the progression of non-canonical forks in which both strands are synthetized by the DNA polymerase delta, which contrasts with the division of labor between DNA polymerase delta and epsilon within origin-born replication forks (Miyabe et al. 2015). Such restarted forks are liable to replication errors such as multiple template switches, replication slippage and U-turn. Despite these unusual features, our data established that a restarted fork remains coupled to histone deposition and thus may help to ensure continuous chromatin assembly upon replication stress.

Fork obstacles and replication stress interfere with the inheritance of epigenetic marks, (Jasencakova et al. 2010; Sarkies et al. 2010; Li et al. 2017). On the one hand, the post-replicative repair of DNA lesions/structures, left un-replicated behind the fork, is uncoupled from chromatin assembly and recycling of parental histones. On the other hand, it was proposed that the bypass of DNA secondary structures, such as G quadruplexes, by PrimPol allows the repriming of DNA synthesis very closly to the fork obstacle and thus reassuring the maintenance of replication-coupled chromatin maturation (Schiavone et al. 2016). The choice of the pathway employed to overcome fork obstacles may impact on the maintenance of histone deposition coupled to restarted forks. Our data indicate that RDR, a main pathway to bypass fork obstacles, is coupled to histone deposition with D-loops offering the possibility to prime repair synthesis-coupled histone deposition. In the view that replisomes are often interrupted by numerous obstacles, RDR-coupled histone deposition contributes to cell resistance to replication stress and may ensure continuity in assembling chromatin upon replication stress.

### Concluding remarks

We propose here that histone deposition coupled to RDR comes at the expense of stabilizing JMs which can be detrimental to genome stability. Overexpression of Asf1b, one isoform of human Asf1, and CAF-1 was found to be associated with poor prognosis in various cancer types (Polo et al. 2010; Corpet et al. 2011). Since cancer cells exhibit signs of replication stress, we speculate that upregulating chromatin assembly factors may favor the stability of at-risk JMs at site of replication stress, channeling RDR events towards a chromosomal rearrangement pathway that fuels cancer development.

## Materials and Methods

### Standards yeast genetics

Yeast strains used in this work are listed in Supplementary Table 1. Gene deletion and gene tagging were performed by classical and molecular genetics techniques (Moreno et al. 1991). Strains containing the *RTS1-RFB* were grown in supplemented EMM-glutamate media containing 60μM of thiamine. To induce the RTS1-RFB, cells were washed twice in water and grown in supplemented EMM-glutamate media containing thiamine (Rtf1 repressed, RFB OFF condition) or not (Rtf1 expressed, RFB ON condition) for 24 hours or 48 hours.

### Chromatin Fraction Assay

Chromatin fractionation was performed as described previously (Kai et al. 2001) with the following modifications. Cells were harvested, digested with 100μg of zymolyase 20T (Amsbio, 120491-1) to obtain spheroplasts, and resuspended in lysis buffer (50mM potassium acetate, 2mM MgCl2, 20mM HEPES pH7.9, 1% Triton X-100, 1mM PMSF, 60mM β-glycerophosphate, 0.2mM Na_3_VO_4_, 1μg/ml AEBSF and Complete Protease Inhibitor EDTA-Free Tablet (Roche, 04 693 159 001). After lysis, extracts were subsequently fractionated into soluble and pellet fractions by centrifugation. The insoluble chromatin-enriched pellet fraction was washed twice with the lysis buffer without 1 % Triton X-100 and digested with 100Units of DNase I HC (ThermoScientific, EN0523) on ice for 15min followed by 15min at RT. The DNase I-digested chromatin-enriched fraction was centrifuged for 5 min at 16,000g. Supernatant was designated as the chromatin fraction. Samples corresponding to total soluble and chromatin fractions were migrated and transferred on nitrocellulose membrane. Proteins of interest were revealed by anti-GFP (1/1000^e^, Roche, 11 814 460 001), anti-MYC (1/300^e^, SantaCruz, 9E10), anti-HA (1/500^e^, SantaCruz, 12CA5), anti-PCNA (1/500^e^, SantaCruz, PC10), antiTubulin (1/4000^e^, Abcam, ab6160) and anti-H3 (1/2000^e^, Abcam, ab1791) antibodies.

### Analysis of replication intermediates by 2DGE

Replication Intermediates (RIs) were analyzed by 2DGE as described in (Ait Saada et al. 2017). RIs were migrated in 0.35% agarose gel in 1X TBE for the first dimension. The second dimension was migrated in 0.9% agarose gel 1XTBE supplemented with EtBr (Brewer 1992). DNA was transferred onto a nylon membrane (Perkin Elmer, NEF988001PK) in 10X SSC. Membranes were incubated with a ^32^P radiolabeled *ura4* probe, and RIs were detected using phosphor-imager software (Typhoon-trio) and quantified with ImageQuantTL.

### Co-immunoprecipitation

5.10^8^ cells were harvested with 0.1% sodium azide, washed in cold water and resuspended in 400 μl of EB buffer (50mM HEPES High salt, 50mM KOAc pH7.5, 5mM EGTA, 1% Triton X-100, 1mM PMSF, and Complete Protease Inhibitor EDTA-Free Tablet (Roche, 04 693 159 001). Cell lysis was performed with a Precellys24 homogenizer (Bertin instruments). The lysate was treated with 250mU/μl of benzonase (Novagen, NOVG 70664-3) for 30min. After centrifugation, the supernatant was recovered and an aliquot of 50 μl was saved as the INPUT control. To 300μl of lysate, 2μl for anti-GFP (Life Technologies, A11122) were added and incubated for 1.5 hours at 4°C on a wheel. Then, 20μl of Dynabeads protein-G (Life Technologies, 10004D) prewashed in PBS were added and then incubated at 4°C overnight. Alternatively, lysates were incubated overnight with 20μl of anti-MYC (Life Technologies, 88842) or anti-HA (Life Technologies, 88836) antibody coupled to magnetic beads. Proteins of interest were detected using anti-HA high affinity (1/500e, Roche, clone 3F10), anti-GFP (1/1000e Roche, 11 814 460 001), anti-MYC (1/300e, SantaCruz, A-14), anti-HA (1/500e, SantaCruz, 12CA5), anti-PCNA (1/500e, SantaCruz, PC10), and anti-H3 (1/2000e, Abcam, ab1791) antibodies.

### Chromatin immunoprecipitation of Pcf1-YFP

ChIP experiments were performed as reported in (Audry et al. 2015). Cells were crosslinked with fresh 1% formaldehyde (Sigma, F-8775) for 15 minutes. Cells were lysed using Precellys24 homogenizer (Bertin instruments) in lysis buffer (50mM Hepes-KOH pH7.5, 140mM NaCl, 1mM EDTA, 1% Triton X-100, 0.1% sodium deoxycholate, 1mM phenylmethylsulfonyl fluoride, and Protease Inhibitor Cocktail (Sigma P8215)). The crude cell lysate was sonicated (using a Diagenod Bioruptor at high setting for 15 cycles: 30 seconds ON + 30 seconds OFF) and clarified by centrifugation for 15min at 16,000g. Prior to immunoprecipitation, 1/100 volume of the cell lysate was saved for an input control. Immunoprecipitations were performed with 2μl of anti-GFP antibody (Life Technologies, A11122) for 2 hours. After 30min incubation with 20μl magnetic beads (Life Technologies, 10004D), immunoprecipitates were successively washed with 2×1ml lysis buffer, 2×1ml lysis buffer/500mM NaCl, 2×1ml wash buffer (10mM Tris-HCl pH8, 0.25M LiCl, 0.5% NP-40, 0.5% sodium deoxycholate, 1mM EDTA) and 1ml TE buffer (10mM Tris-HCl, 1mM EDTA pH8). Crosslink was reversed by incubating the samples at 65°C overnight. Samples were then treated with 0.5mg/ml Proteinase K (Euromedex, EU0090) and DNA was purified using Qiagen PCR purification kit and eluted in 100μl of water. The relative amount of DNA was quantified by qPCR (primers are listed in supplementary Table 2). Pcf1-YFP enrichment was normalized to an internal control locus *(ade6)*.

### Pulse-Field Gel Electrophoresis

PFGE were performed as described in (Lambert et al. 2010). Membranes were then incubated with a ^32^P radiolabeled *rng3* probe. Quantification of acentric chromosomes visualized by PFGE was performed as described in (Pietrobon et al. 2014).

### Microccocal digestion and BrdU incorporation

Microccocal digestions were performed as described in (Pai et al. 2014). After crosslink with 1% formaldehyde, 1.10^9^ Cells were spheroplasted in 2ml of CES buffer (50mM Citric acid/50mM Na2HPO_4_ pH 5.6, 40mM EDTA pH8, 1.2M Sorbitol, 20mM β-mercaptoethanol) containing 1 mg/ml Zymolyase 100T (Amsbio, 120493-1) for 20min at 30°C. Spheroplasts were washed twice with 1ml of iced cold 1.2M Sorbitol buffer. Cells were resuspended in 1ml of NP-S buffer (1.2M Sorbitol; 10mM CaCl_2_, 100mM NaCl, 1mM EDTA pH8, 14mM β -mercaptoethanol, 50mM Tris-HCl pH8, 0.075% NP-40, 5mM spermidine, 0.1mM PMSF, Complete Protease Inhibitor EDTA-Free Tablet (Roche, 04 693 159 001)) containing the indicated units of Micrococcal Nuclease (Worthington Biochemical, LS004798) for 10 min at 37°C. Reactions were stopped by addition of 50mM EDTA pH8 and SDS 0,2%. Crosslinking was reversed overnight at 65°C in the presence of 20μg of RNAseA (Sigma, R5503) and 0.2mg/ml Proteinase K (Euromedex, EU0090). DNA was purified by phenol/chloroform extraction and ethanol precipitation. Purified DNA was resolved on 1.5% agarose gel (1X TBE).

For BrdU incorporation, we used cells expressing *Drosophila melanogaster* deoxyribonucleoside kinase (DmdNK) under the control of the fission yeast adh promoter, together with the human equilibrative nucleoside transporter (hENT1) *(adh-dmdNK-adh-hENT1)* (Fleck et al. 2017). Cells were arrested 4 hours with 40mM HU (Sigma, H8627) and released in fresh media containing BrdU (Sigma, B5002) for 20min. After MNase digestion, DNA was analyzed by Southern-blot using anti BrdU antibody (1/4000^e^, Abcam, ab12219).

### Spontaneous Recombination assay

Spontaneous recombination rate was assayed using strains containing a direct repeat of two nonfunctional *ade6* alleles flanking a functional *ura4* gene (Hartsuiker et al. 2001). Strains were kept on low adenine EMM plates lacking uracil to prevent selection for Ade^+^ and Ura^-^ recombinants. Dark pink colonies were streaked on supplemented YE plates and at least 11 independent single colonies for each strain were used to calculate Ade+ recombinant rate. Appropriate dilutions were plated on supplemented YE plates (to determine cell survival), EMM plates lacking adenine (to score spontaneous recombination rate, Ade+ recombinants) and EMM plates lacking adenine and uracil (to score gene conversion rate, Ade+ Ura+ recombinants). Colonies were counted after 5-7 days of incubation at 30°C.The rates of Ade+ and Ade+ Ura+ recombinant were calculated as described in (Lea and Coulson 1949).

### Analyzed X-ray structures of nucleosome

There is no experimental model of nucleosome containing histones from *S. pombe*. Among the numerous available structures of nucleosome, only three of them include yeast histones, all from *S. cerevisiae* fPDB codes 1ID3, 4JJN and 4KUD), with resolutions of ~3 Å. Indeed, the highest resolution was obtained for a nucleosome containing histones from *Xenopus laevis* (PDB code 1KX5, resolution of 1.9 Å). Since the histone sequences are extremely conserved, *S. cerevisiae* and *Xenopus laevis* H3 histones share 92% of residues with H3 of *S. pombe*. More specifically, the region surrounding H113 is well preserved across *S. pombe, S. cerevisiae* and *Xenopus laevis*, with a very good score being observed for the couple *S. pombe / Xenopus laevis* (Figure S2A). Given the reasonable sequence agreement, 1ID3, 4JJN, 4KUD and 1KX5 were analyzed using PDBsum (de Beer *et al*, 2014) to describe the interface between two H3-H4 dimers (de Beer et al. 2014).

## Acknowledgements

We thank Makoto Kawamukai for the gift of the *asf1-myc* and *asf1-33-myc* strains and Edgar Hartsuiker for the gift of the *adh-dmdNK-adh-hENT1 strain*. We thank Jean-Pierre Quivy, Vincent Pennaneach and Kirill Lobachev for critical comments of the manuscript. We also thank the PICT-IBiSA@Orsay Imaging Facility of the Institut Curie. This study was supported by grants from the Institut Curie, the CNRS, the *Fondation ARC*, the *Fondation Ligue (comité Essonne), l’Agence Nationale de la Recherche* ANR-14-CE10-0010-01, the *Institut National du Cancer* 2016-1-PLBIO-03-ICR-1 and the *Fondation pour la Recherche Médicale* “Equipe FRM DEQ20160334889”. ATS and AAS were funded by the Institut Curie international PhD program, and a French governmental fellowship, respectively. ATS was supported by a 4^th^ year PhD fellowship from *Fondation pour la Recherche Medicale* (FDT20160435131). The funders had no role in study design, data collection and analysis, the decision to publish, or preparation of the manuscript.

## Author contributions

J.H., D.D., A.A.S., A.T.S., L.D., F.M. and K.F. performed the experiments. B.H. and F.O. provided structural biology expertise. J.H. and S.A.E.L. designed experiments and analyzed the data. J.H. and S.A.E.L. wrote the paper.

## Competing financial interests

The authors declare no competing financial interests.

